# Cavity-nesting bees combine forest nesting habitats with surrounding floral resources in a subtropical forest diversity experiment

**DOI:** 10.64898/2026.05.20.726496

**Authors:** Ting-Ting Zhang, Massimo Martini, Juan-Juan Yang, Guo-Ai Chen, Huan-Xi Cao, Qing-Yi Yu, Finn Rehling, Ming-Qiang Wang, Michael C. Orr, Manuela Sann, Felix Fornoff, Jing-Ting Chen, Qing-Song Zhou, Ze-Qing Niu, Christina M. Grozinger, Xiao-Juan Liu, Alexandra-Maria Klein, Chao-Dong Zhu, Arong Luo

## Abstract

Wild bees face declines, and forests may serve as critical habitats for pollinators. However, how forest composition and the associated floral environment shape pollen provisioning and resource partitioning among cavity-nesting bees remains poorly understood. Here, we leveraged BEF–China, a large-scale subtropical forest biodiversity experiment with experimentally controlled plant (tree and shrub) communities, to investigate how forest composition and spatial context shape pollen provisioning, resource partitioning, and reproductive success of cavity-nesting bees.

We used DNA metabarcoding to analyze floral composition of pollen provisioned by five cavity-nesting bee species, with samples collected from BEF–China across three years (2022– 2024). By comparing pollen taxonomic composition from whole-nest pooled samples and individual brood-cell samples with the experimentally planted species pool, we characterized dietary patterns and temporal dynamics of five bee species.

Bees primarily relied on floral resources from the surrounding landscape, with planted trees providing essential but temporally restricted pollen supplements during specific phenological stages. Co-occurring bee species exhibited staggered nesting phenology and distinct dietary preferences for different plant families, with fine-scale resource differentiation even during periods of phenological overlap.

Our results suggest that managed forests support cavity-nesting bees by providing critical woody floral resources during specific phenological gaps and offering stable nesting environments. To mitigate pollinator declines, forest management should prioritize maintaining diverse, phenologically complementary flowering vegetation within and surrounding forest stands. This ensures temporal continuity of pollen availability throughout the nesting season, which is particularly crucial for restoring pollinator services in simplified forest landscapes.

## Introduction

Bees are essential pollinators in both natural and agricultural ecosystems, supporting roughly 87.5% of angiosperms and two-thirds of major food crops (Klein et al., 2007; Luo et al., 2026; Ollerton et al., 2011), with solitary species accounting for approximately 90% of the more than 20,000 species globally (Ascher and Pickering, 2025). However, wild bee populations are experiencing widespread declines globally due to habitat loss, intensive agriculture, pesticide exposure, and climate change (Guzman et al., 2024). Forests cover approximately 31% of the global land area and harbor a substantial proportion of terrestrial biodiversity (Brockerhoff et al., 2017; FAO, 2020). These ecosystems may serve as critical habitats for bee conservation in several ways. Forests provide nesting substrates (e.g., deadwood, hollow stems) and diverse floral resources across extended flowering seasons, while buffering microclimate extremes that could influence bee development (Eckerter et al., 2021; Ulyshen et al., 2023). Understanding how forest structure and composition support bee communities is needed to incorporate pollinator needs into forest management. This is particularly critical for solitary bees, as the availability and diversity of floral resources directly influence their ability to provision their offspring with adequate pollen-nectar mixtures during their 4-8 week adult lifespan (Danforth et al., 2019; Vaudo et al., 2018).

Biodiversity–ecosystem functioning (BEF) theory predicts that plant species richness enhances consumer diversity through bottom-up pathways. This prediction has been supported by studies on arthropod functional groups (Haddad et al., 2011; Schuldt et al., 2019) and their multitrophic interactions in experimental settings (Cao et al., 2018; Li et al., 2025; Scherber et al., 2010). In a grassland biodiversity experiment, brood cell density of cavity-nesting bees increased with flowering plant diversity (Ebeling et al., 2012). However, this pattern was not observed in a subtropical forest biodiversity experiment, where tree diversity showed limited association with bee diversity (Guo et al., 2021). This discrepancy may reflect fundamental differences in how grassland and forest ecosystems support bee communities. Limited understory flora in closed-canopy forests, combined with foraging ranges (150–600 m) (Gathmann and Tscharntke, 2002) surpassing plot dimensions, complicates local-scale resource assessments for solitary bees. The relative importance of floral resources from within experimental plots versus the surrounding habitat matrix for bee provisioning remains poorly quantified, and the role of forest floral resources in shaping cavity-nesting bee assemblages is still unclear.

Beyond floral resource distribution, the spatial position of nest within a forest stand may independently affect bee reproductivity. Forest interiors buffer microclimate extremes and may reduce parasitoid pressure, benefiting offspring development (De Frenne et al., 2019; Zellweger et al., 2020), whereas forest edges typically support higher floral resource density due to greater light availability (Mullally et al., 2019; Ren et al., 2023). As obligate central-place foragers, cavity-nesting bees are spatially constrained by the necessity of returning to a fixed nest site to provision their offspring (Woodgate and Chittka, 2018). This potential trade-off between nesting environment quality and foraging efficiency means that the net effect of nest site selection on cavity-nesting bee reproduction in managed forests remains an open question. Standardized trap-nest directly links bee foraging to reproduction by allowing for the simultaneous measurement of resource use and nesting success (Staab et al., 2018; Tscharntke et al., 1998). With the aid of DNA metabarcoding, pollen samples provide high-throughput, taxonomically resolved information on floral diet (Encinas-Viso et al., 2023; McFrederick and Rehan, 2016). The BEF–China platform provides a suitable system for applying these methods. The platform experimentally manipulates tree species composition across a large number of plots and has now reached a mature stage, planted vegetation flowers regularly, canopy closure is established, and ground-layer vegetation is removed annually. Embedded within a mosaic of surrounding habitats, the platform is accessible to solitary bees whose foraging ranges extend beyond individual plots, allowing potential integration of floral resources from multiple habitat types. By comparing pollen taxonomic composition with experimentally planted forest stand species pool, these methods enable empirical assessment of the extent to which bees provision offspring from forest plant communities versus vegetation in surrounding heterogeneous matrices.

The dominant cavity-nesting bees on the platform are megachilid bees (Anthophila: Megachilidae), with five species regularly occupying trap nests during the main flowering season, as established by prior monitoring on the same platform (Fornoff et al., 2021). These species vary in body size and nesting phenology, offering natural variation for investigating resource partitioning. Here, we combined three years of trap nest monitoring (April–October, 2022–2024) with DNA metabarcoding of 412 pollen samples from five co-occurring bee species, using both whole-nest pooling and individual brood-cell sampling, to address three questions: (1) What floral resources do cavity-nesting bees provision to offspring in closed-canopy forests, and do these resources primarily originate from within experimental plots or from the surrounding habitat matrix? (2) Do co-occurring species partition resources temporally or dietarily? (3) How does the edge positioning of experimental plots contribute to bee provisioning patterns and reproductive capacity?

We predicted that (1) pollen provisions would be dominated by planted species if experimental trees provided sufficient floral resources during bee nesting periods, but would shift toward surrounding vegetation if plot-level floral resources were limited or phenologically mismatched; (2) co-occurring species would reduce competition through staggered phenology or dietary specialization; and (3) at the scale of their foraging range, nest productivity would be higher in forest interior plots even if forest edges provide more accessible floral resources.

## Material and Methods

### Study site and experimental design

This study was conducted on the BEF–China experimental forest, located in Xingangshan, Jiangxi Province, China (29.08°–29.11°N, 117.90°–117.93°E). The area is characterized by a subtropical monsoon climate with a mean annual temperature of 16.7 °C and average precipitation of 1821 mm. The BEF–China platform comprises 566 plots (25.8×25.8 m) across two sites (A and B), established in 2009 and 2010, respectively (Bruelheide et al., 2014). The surrounding area includes forests, agricultural land, and early successional vegetation. Tree species richness was experimentally manipulated from monoculture to high-diversity mixtures (400 trees per plot), with plots containing balanced mixtures of evergreen and deciduous species embedded within surrounding forest matrix. By the time of this study, canopy closure was largely complete (Deng et al., 2025), and most planted trees had reached flowering age, as evidenced by regular seed production and phenological records compiled from regional floras and climate data (BEF–China management team, unpublished data). Understory vegetation within experimental plots is removed yearly to maintain the original tree richness gradient, resulting in minimal flower resources in the forest floor.

### Trap nest deployment and bee sampling

We used pollen samples collected from 2022 to 2024. In 2022–2023, reed-based trap nests were deployed following the study by (Fornoff et al., 2021). Two centrally positioned poles were placed 11 m apart in each of 88 plots covering all levels of tree species richness (Table S1), with each pole supporting two standardized artificial trap nests. Occupied reed tubes were collected monthly from April to October and transported to the laboratory, where nests were reared individually in glass tubes at ambient temperature until adult emergence (Staab et al., 2016). All emerged bees were identified morphologically to species level, with reference to a voucher collection of trap-nest reared specimens established by C-D Zhu’s lab (Institute of Zoology, Chinese Academy of Sciences, Beijing).

In 2024, layer trap nests consisting of stacked medium-density fiber (MDF) boards with pre-drilled cavities of varying diameters (2–15 mm) were used to attract cavity-nesting bees. These traps were designed to allow direct access to individual brood cell contents during the nest construction period. One layer nest was placed at the center of each plot on a wooden pole at around 1.3 m height. Each nest comprised 14 layers (15×15 cm, 1.9 cm thick) with seven cavities per layer, providing 98 total nesting cavities (Klein et al., 2026). Layers were secured with a tension strap and protected from rain with a plastic cover. Traps were exposed from late April to mid-August, and bee species were identified based on nesting biology characteristics and DNA barcoding of the cytochrome C oxidase subunit I (COI) gene region, using molecular reference databases (Wang et al., 2025) to achieve species-level identification without rearing.

### Pollen sampling strategies

Two complementary pollen sampling approaches were employed between years and trap type: pooling pollen from all brood cells within each individual nest in 2022–2023 and sampling individual brood cells separately in 2024. For whole-nest pooling, contents of all brood cells within each constructed nest were pooled in equal proportions to generate a single composite sample per nest (Fig. 1a), providing species-level estimates of overall foraging breadth and dietary composition. For individual-cell sampling, pollen was collected randomly from up to three brood cells separately as independent samples from a single tube (Fig. 1b), enabling assessment of within-nest variation in pollen provisioning. Pollen samples were stored at -80 °C until DNA extraction.

**Figure 1.**
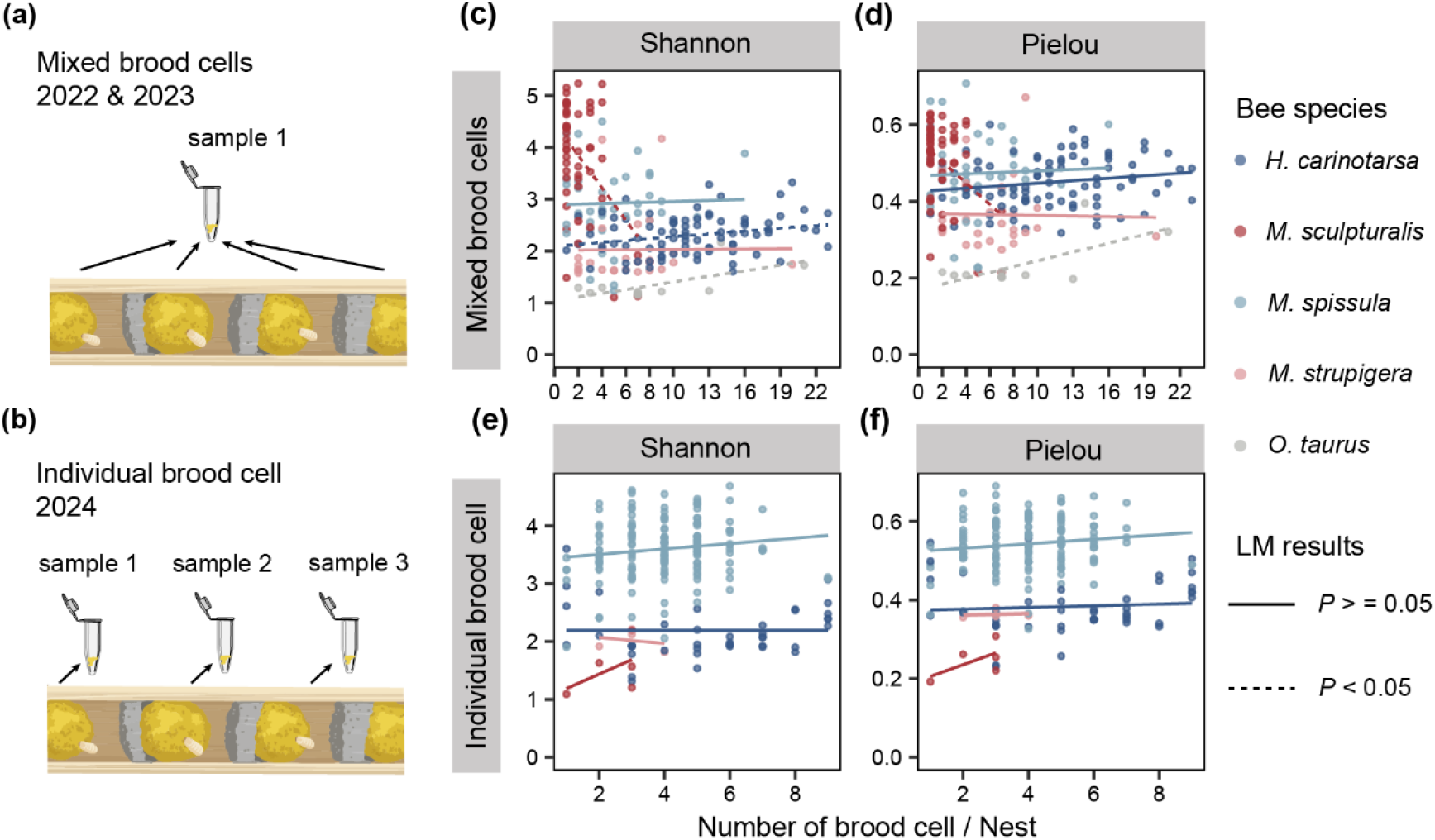
Relationship between pollen diversity and brood cell number per nest across solitary bee species. Pollen diversity (Shannon index, c, d) and evenness (Pielou index, e, f) were calculated from two sampling strategies: mixed pollen samples from all brood cells within a nest (a) and individual brood cell pollen samples (b). Each point represents a pollen sample for sequencing. Different colors indicate different bee species as shown in the legend. Solid lines represent model-predicted trends, and dashed lines indicate statistically significant relationships (*P* < 0.05) based on LMs. See Table S4 for full statistical analyses controlling for spatiotemporal factors.

### DNA extraction and amplification

Total genomic DNA (gDNA) was extracted from homogenized pollen samples using a modified CTAB protocol (Cullings, 1992). The plastid *rbcL* barcode region was amplified using primers rbcLaF and rbcLaR (Erickson et al., 2017; Kress and Erickson, 2007), and sequenced on an Illumina NovaSeq PE250 platform (see Supplementary Methods S1 for details).

### Bioinformatics and taxonomic assignment

Raw sequences were quality-filtered, denoised into zero-radius operational taxonomic units (ZOTUs) using USEARCH v10 (Edgar, 2013) and the UNOISE3 algorithm (Edgar, 2016), and taxonomically assigned against a curated *rbcL* reference database (Bell et al., 2017; Weber et al., 2023). After taxonomic aggregation, ZOTUs were assigned to 54 families and 76 genera. All datasets were rarefied to 17,461 sequences per sample, corresponding to the lowest number of sequences observed among all samples, to ensure equal sampling effort prior to diversity comparisons (see Supplementary Methods S2 for full bioinformatics pipeline).

### Statistical analyses

All statistical analyses were performed in R v4.2.2. We focused on the target megachilid species that dominated trap-nest occupancy during 2022–2023 (*Hoplitis carinotarsa* Wu, *Megachile sculpturalis* Smith, *Megachile spissula* Cockerell, *Megachile strupigera* Cockerell, and *Osmia taurus* Smith) and counted their monthly nest abundance (log2-transformed within species). Regional flowering phenology was estimated from local flora and climate records (compiled by Prof. Bo Yang, BEF–China), showing monthly counts of flowering species including pollen-producing gymnosperms. To ensure statistical robustness and reduce stochastic variation, we retained only bee species with at least three pollen samples per month per year, resulting in a final dataset of 412 samples, including 83 from 2022, 129 from 2023, and 200 from 2024 (Table S2). To assess the consistency between the two sampling approaches, we assessed correlations between pollen diversity (Shannon and Pielou indices) and the number of brood cells per nest using linear models (LMs). For the 2022–2023 dataset, we further applied linear mixed-effects models (LMMs) or LMs to account for spatiotemporal variation, with sampling plot included as a random intercept and month and year as fixed effects (Bates et al., 2015). We further compared the within-nest variation revealed by the 2024 individual-brood-cell sampling approach, quantifying differences in pollen composition among cells using Morisita–Horn dissimilarity and examining variation in the identity and relative abundance of dominant plant genera. Because individual brood cell samples showed substantial within-nest variation, subsequent community-level analyses utilized exclusively whole-nest pooled samples from 2022–2023 to maintain temporal consistency across years.

To assess the origin of pollen resources provisioned by cavity-nesting bees, we calculated the proportion of sequence reads corresponding to plant genera not present in experimental plantings as an indicator of extra-plot foraging. We selected pollen genera contributing >97% of mean sequence reads across the four focal bee species (*O. taurus* excluded due to limited resolution of Fagaceae by the *rbcL* marker), then classified them as woody or herbaceous based on plant growth form. Both overall resource use patterns and monthly genus-level composition for each species were visualized.

To characterize temporal foraging dynamics and dietary differentiation among co-occurring species, we calculated monthly mean Shannon diversity and mean brood cell production for each bee species. We quantified the magnitude of seasonal shifts in foraging composition using centroid displacement, and tested for community differences in Morisita–Horn dissimilarity using PERMANOVA (Bramon Mora et al., 2020; Legendre, 2014). Interannual consistency in pollen composition was visualized using stacked histograms, which showed monthly pollen family composition for each bee species, with the eight most abundant families displayed separately and the remaining low-abundance families combined as “others” (Milla et al., 2022; Sickel et al., 2015).

To evaluate the effects of forest composition and spatial position on bee nesting patterns, we first tested whether bee assemblage and species-level responses (assemblage nest abundance, assemblage species richness, and species-specific nest abundances) were related to tree species richness using generalized linear mixed models (GLMMs) implemented in the glmmTMB package (Brooks et al., 2017). We then assessed the effects of nest position relative to the platform edge. Using 5-m resolution shapefiles of Site A and Site B boundaries (WGS 84 coordinate system), we calculated the nearest distance from each of the 88 trap-nest plots to the respective site edge. We tested the effects of distance to edge on both nest abundance and nest productivity using GLMs or GLMMs depending on data structure. For nest abundance (plot-level), models included distance, collection year, and site as fixed effects. For nest productivity (individual nest-level, measured as number of brood cells per nest), we excluded *O. taurus* due to insufficient sample size and limited temporal coverage. We tested for distance × year interactions to assess whether edge effects varied between years; as these were non-significant, we pooled data across years and included sampling month as a covariate. To minimize confounding between temporal and spatial patterns, analyses focused on months of peak nesting activity for each species. Poisson or negative binomial error distributions were used depending on overdispersion diagnostics (Lüdecke et al., 2021). All model diagnostics, including checks for residual patterns and overdispersion, were performed using the DHARMa package (Hartig, 2024).

## Results

### Nest abundance patterns and pollen sequencing results

Across 2022–2023, a total of 912 occupied nests were recorded for the five focal species. No significant relationship was detected between local plot tree species richness and bee assemblage nest abundance (GLMM: estimate = -0.104, *P* = 0.197) or bee assemblage species richness (GLM: estimate = -0.009, *P* = 0.762; Table S3). However, species-specific analyses revealed divergent responses that masked patterns at the assemblage level. Specifically, the abundance of *H. carinotarsa* declined significantly with increasing tree species richness (GLMM: estimate = -0.357, *P* = 0.003; Table S3). The remaining species showed variable responses to tree richness, with trends varying in direction (both positive and negative) but lacking statistical significance (Table S3). Model diagnostics (DHARMa residuals) confirmed adequate model fit for all species-level analyses (Fig. S1). Bee species exhibited distinct nesting phenology, with activity peaks varying among species and between years (Fig. S2). Estimated BEF–China platform flowering phenology showed floral resources were available throughout the trap-nest monitoring period, with highest diversity in spring months (Fig. S3).

A total of 412 pollen samples yielded 23,083,935 sequences, with individual sample counts ranging from 17,461 to 155,200 reads. Following quality filtering, denoising, and taxonomic assignment, 1,143 ZOTUs corresponding to common flowering plant taxa were identified and retained, with sequence lengths ranging from 326 to 367 bp. After taxonomic aggregation, these ZOTUs represented 54 families and 76 genera. Rarefaction curves for all five bee species reached asymptotes, confirming adequate sequencing depth (Fig. S4a). The rank-abundance distribution exhibited a steep curve with a long tail, indicating that bee foraging was concentrated on a small number of dominant ZOTUs, with numerous low-abundance ZOTUs representing only minor dietary components (Fig. S4b).

### Comparison of pollen sampling methods

Pollen diversity (Shannon and Pielou indices) showed species-specific relationships with brood cell number per nest (Fig. 1c-f; Table S4). *M. sculpturalis* showed a significant negative relationship (Shannon: estimate = -0.092, *P* = 0.001; Pielou: estimate = -0.009, *P* = 0.004; Table S4), while other species exhibited positive trends of varying significance (Table S4). Analysis of within-nest variation using Morisita–Horn dissimilarity revealed substantial heterogeneity among brood cells, with some nests exhibiting highly similar pollen compositions and others differing markedly (Fig. S5a, S5b). Subsequent analyses focused on mixed-brood-cell samples, which provide more representative species-level foraging patterns than individual cells and enable linkage to nest-level reproductive outcomes.

### Seasonal dynamics of pollen diversity and reproductive output

Pollen diversity, nest productivity, and pollen community composition had distinct temporal variation among the five bee species (Fig. 2). For *M. spissula* and *M. sculpturalis*, an inverse relationship emerged between pollen diversity and brood cell counts, with months of higher reproductive output corresponding to lower pollen diversity (Fig. 2a, 2b). This pattern was less pronounced in *M. strupigera* and *H. carinotarsa* (Fig. 2a, 2b). Quantification of temporal stability analysis revealed contrasting patterns among species. *M. strupigera* and *H. carinotarsa* exhibited relatively stable dietary compositions across months (Fig. 2c; Table S5). In contrast, *M. spissula* and *M. sculpturalis* showed significant dietary shifts during specific periods, particularly from August to September (Fig. 2c; Table S5).

**Figure 2.**
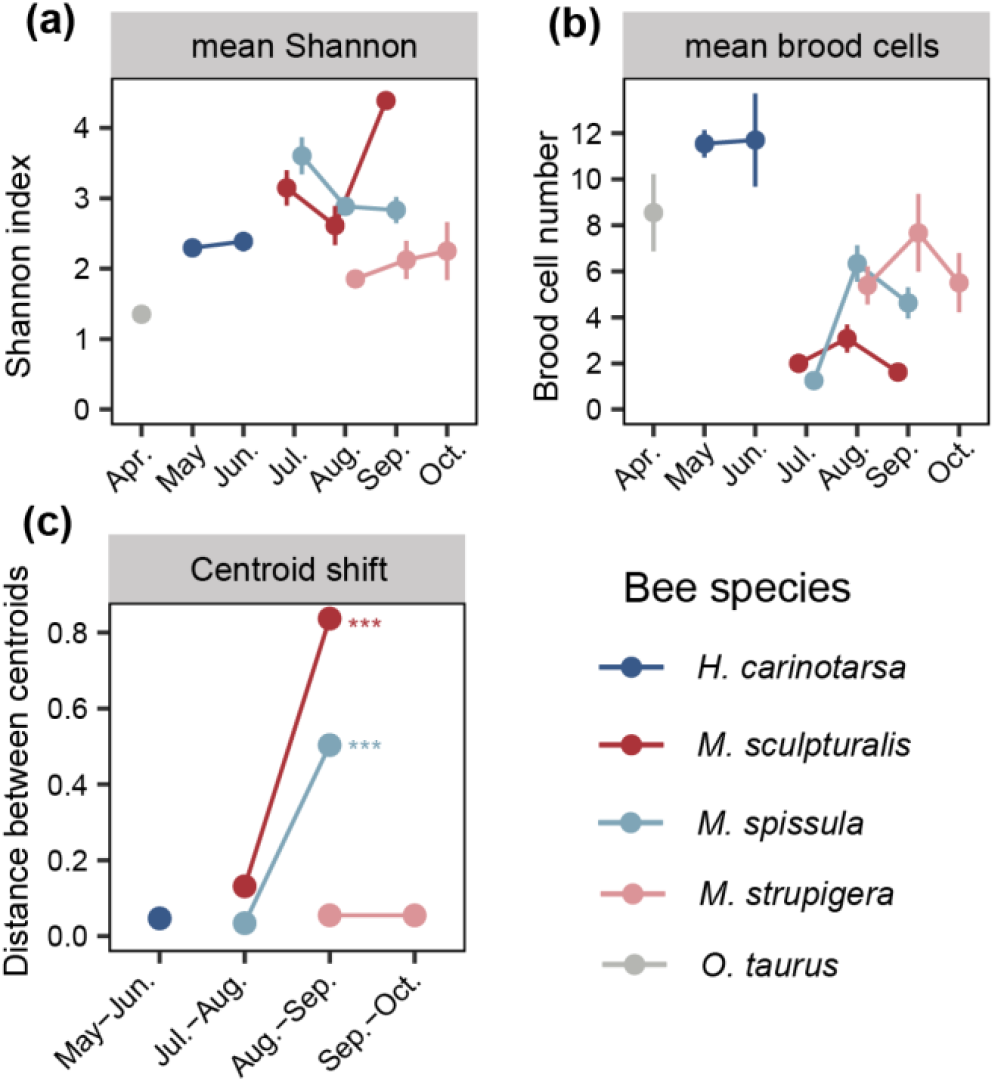
Seasonal variations in pollen diversity, nest productivity, and inter-month community shifts among solitary bee species. (a) Mean Shannon diversity index of pollen collected by each bee species across different months. (b) Mean number of brood cells per nest for each bee species across different sampling months, reflecting nest productivity. Error bars in (a) and (b) represent the standard error of the means. (c) Centroid shift calculated using Morisita–Horn distance, showing the dissimilarity in pollen community composition between adjacent months for each bee species. Significance was tested using PERMANOVA (*** *P* < 0.001; see Table S5 for full details).

### Interspecific resource partitioning and extra-plot pollen sources

Across the remaining four focal bee species, on average, 89.49% of sequence reads derived from plant genera absent from experimental plantings (mean of species-level averages; range: 64.36–99.96% across species; Table S6), confirming that bees foraged predominantly beyond plot boundaries. This pattern was most pronounced from May to August, coinciding with peak nesting activity (Table S6).

Pollen family composition differed markedly among bee species, particularly during peak flowering months (July–September; Fig. 3a, Fig. S6-7), reflecting both species-specific floral preferences and staggered nesting phenology. In August, when multiple species were simultaneously active, dietary segregation was pronounced, with *M. spissula, M. sculpturalis*, and *M. strupigera* foraged primarily on Rutaceae, Fabaceae, and Lamiaceae, respectively, indicating stable species-specific preferences at these sites (Fig. 3a). Genus-level analysis of dominant pollen taxa (collectively accounting for >97% of sequence reads) further revealed differentiation in woody versus herbaceous plant use (Fig. 3b). *H. carinotarsa* predominantly foraged on woody *Ilex* and herbaceous *Uncaria, M. sculpturalis* favored Fabaceae genera (woody *Styphnolobium* and herbaceous *Pueraria*), *M. strupigera* showed strong preference for herbaceous *Vitex*, while *M. spissula* utilized woody *Tetradium, Rhus*, and *Koelreuteria* (Fig. 3b). Notably, the dominant pollen genera across the majority of species originated from non-experimental vegetation (Fig. 3b, Fig S7b).

**Figure 3.**
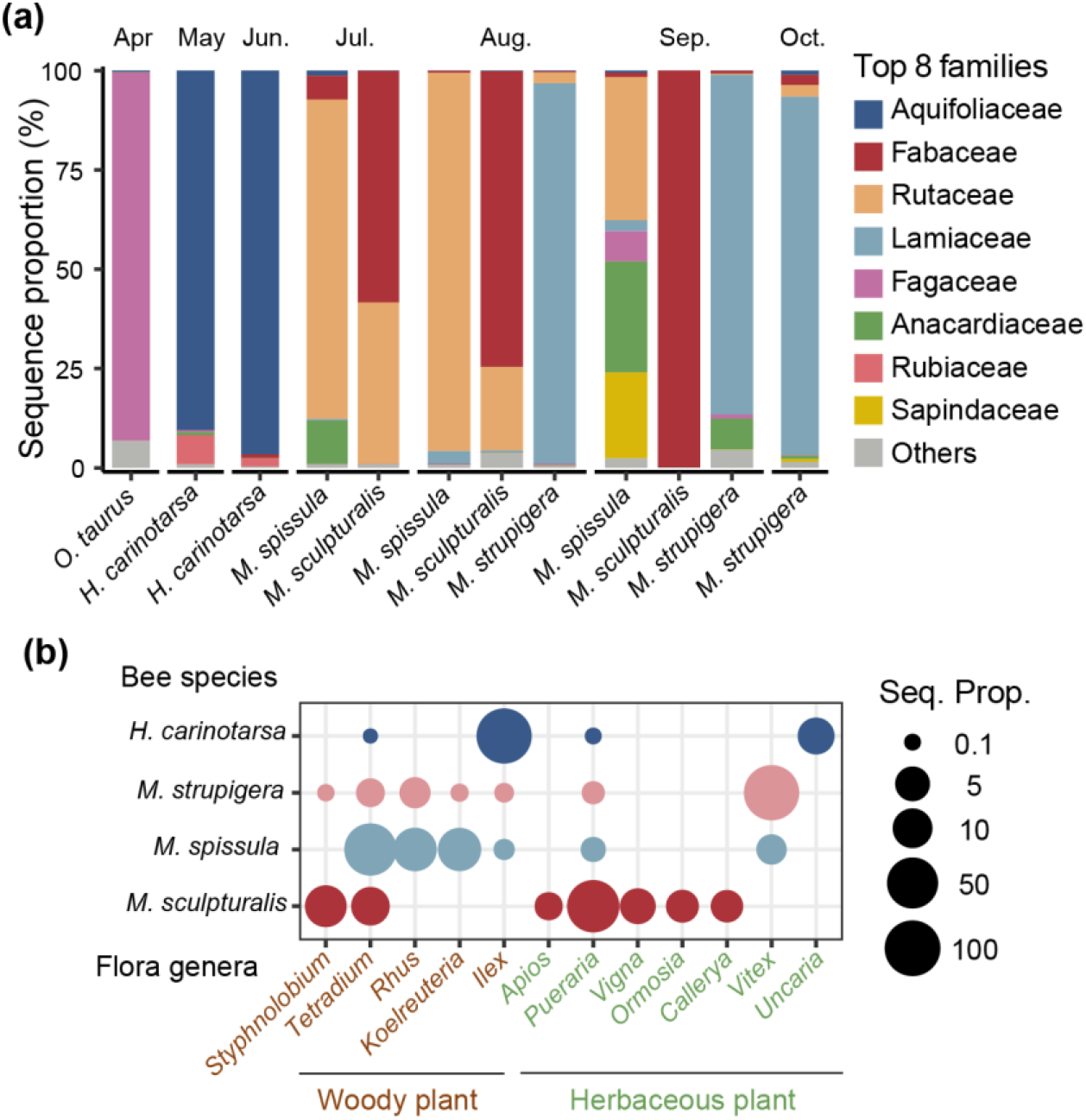
Temporal pollen composition of cavity-nesting bees. (a) Sequence proportions of the top-eight pollen families collected by different solitary bee species in 2022 and 2023, showing monthly compositional shifts. Within each bar, different colors represent the pollen families, while the colored bar at the bottom of each stack indicates the collection month. (b) Sequence proportions of the top 10 dominant pollen genera for four selected bee species. The size of each circle corresponds to the log2-transformed relative proportion of sequencing reads. The log2 transformation was applied to improve visual discrimination among genera with large differences in relative abundance. Plant growth forms were classified into woody plants and herbaceous plants.

### Effects of distance to platform edge on nesting patterns

Distance to platform edge showed different patterns for nest abundance and nest productivity. For nest abundance, no species exhibited significant distance effects (all *P* > 0.08; Table 1). In contrast, nest productivity (brood cells per nest) increased significantly with distance to edge for *M. spissula* (estimate = 0.199, *P* = 0.042), while *H. carinotarsa, M. sculpturalis*, and *M. strupigera* showed no significant distance-fecundity relationships (all *P* > 0.11; Table 1, Fig. S8), although the direction of estimated effects was consistently positive.

**Table 1.** Effects of distance to platform edge on nest abundance and nest productivity of cavity-nesting bee species.

| Bee species | Nest abundance |  |  | Nest productivity |  |  |
| --- | --- | --- | --- | --- | --- | --- |
|  | n | Estimate | *P* | n | Estimate | *P* |
| *H. carinotarsa* | 351 | 0.007 | 0.333 | 322 | 0.059 | 0.223 |
| *M. sculpturalis* | 121 | -0.013 | 0.084 | 109 | 0.139 | 0.111 |
| *M. spissula* | 216 | -0.001 | 0.959 | 186 | 0.119 | **0.042** |
| *M. strupigera* | 191 | -0.005 | 0.507 | 170 | 0.098 | 0.175 |
| *O. taurus* | 35 | 0.025 | 0.195 |  |  |  |

## Discussion

Cavity-nesting bees in experimental forest stands face a spatial challenge, as nesting and foraging resources are distributed across multiple habitat types. Our results show that bees integrate forest nesting habitats with floral resources from the surrounding landscape, with planted trees providing phenologically restricted but important pollen supplements such as Fagaceae in early spring and Anacardiaceae in autumn. Because foraging ranges of these species (150–600 m) extend well beyond individual plot dimensions, this cross-habitat provisioning obscures relationships between plot-level tree diversity and bee abundance. Co-occurring species further exhibited distinct dietary preferences and staggered nesting phenology, reducing competitive overlap. Nest productivity was positively associated with distance from platform edges, suggesting that forest interior conditions benefit offspring development.

### Pollen resources from the surrounding matrix supplemented the forest flower resources

As expected, planted trees contributed to pollen supply, but only during specific phenological windows, such as *O. taurus* foraged for Fagaceae pollen in early spring, and *Rhus chinensis* and *Koelreuteria bipinnata* served as important autumn pollen sources for *M. spissula*. Beyond these windows, the majority of pollen was derived from non-experimental vegetation, indicating that plot-level floral resources alone were insufficient to sustain bee provisioning throughout the nesting season.

Because most provisioned pollen originated beyond plot boundaries, tree species richness within a 25.8×25.8 m plot had limited influence on what bee provision to offspring. In cavity-nesting bees, females select nest sites based on microhabitat cues such as cavity dimensions and substrate conditions rather than surrounding floral composition (Danforth et al., 2019), and subsequently acquire pollen across a broader foraging range (Zurbuchen et al., 2010). In closed-canopy plantation forests, annual management removes understory vegetation and canopy closure limits light availability to the forest floor, leaving the ground layer dominated by ferns and thatch grass that provide minimal, if any, floral resources for bees (Xie et al., 2026). Pollen provisioned by cavity-nesting bees was therefore supplemented extensively by the surrounding matrix rather than confined to experimentally planted species, breaking the link between local tree diversity and bee provisioning. This pattern differs from grassland biodiversity experiments, where plant diversity positively predicted brood cell density because floral resources and nesting substrates co-occurred within bee foraging ranges (Ebeling et al., 2012).

### Temporal and dietary differentiation among co-occurring species

Co-occurring bee species partitioned floral resources through both dietary specialization and temporal segregation (Ye et al., 2024; Zaragoza-Trello et al., 2023). During peak flowering periods, species showed distinct preferences for different plant families, such as *M. spissula* on Rutaceae, *M. sculpturalis* on Fabaceae, and *M. strupigera* on Lamiaceae, reducing direct dietary overlap. Staggered nesting phenology further minimized competition, with species concentrating reproductive effort at different times. Species also varied in their temporal foraging stability, with *M. strupigera* and *H. carinotarsa* maintaining consistent diets while *M. spissula* and *M. sculpturalis* shifted diet composition tracking flowering phenology (Fig. 2c, Table S5, Fig. S7).

The inverse relationship between monthly pollen diversity and brood cell production in *M. spissula* and *M. sculpturalis* (Fig. 2a, 2b) suggests that high reproductive output depends on access to abundant, concentrated floral resources rather than a broad diet (Vaudo et al., 2016). This pattern may be particularly pronounced in large-bodied species such as *M. sculpturalis*, which can exploit mass-flowering events within their greater foraging range. Provisioning trade-offs between pollen diversity and quantity may be further related to the nutritional quality of resources as well as offspring traits such as sex and body size (Kapheim et al., 2011; Vaudo et al., 2024).

### Forest interior conditions and tree diversity effects on reproductive success

Tree species richness showed no significant association with plot-level bee abundance or species richness at the assemblage level (Table S3), consistent with previous findings from the same platform (Guo et al., 2021). The exception was *H. carinotarsa*, which declined significantly with increasing tree richness, possibly reflecting species-specific sensitivity to microhabitat conditions that differ between monoculture and mixed stands. This divergence shows that assemblage-level patterns can mask biologically meaningful responses among individual species.

Nest productivity increased with distance from the platform edge in *M. spissula*, with positive but non-significant trends in the remaining species (Table 1). This contrasts with the expectation that forest edges, being adjacent to open habitats with higher floral resource density (Eckerter et al., 2022), would benefit reproduction by reducing foraging costs (Stuligross and Williams, 2020). That nest productivity was not higher near edges despite this potential advantage suggests that edge-associated costs, such as greater parasitoid pressure (Steffan-Dewenter, 2002; Tylianakis et al., 2007) or less stable nesting conditions (Forrest and Chisholm, 2017; Vincze et al., 2024), outweigh the benefits of floral proximity. The positive edge-distance effect thus likely reflects a trade-off between foraging accessibility and nesting environment quality, although our data cannot resolve which specific mechanisms drive this pattern.

### Methodological considerations

Our study employed two complementary sampling strategies to capture different aspects of foraging behavior. Whole-nest pooling provided a robust overview of species-level dietary composition, but individual-cell analysis revealed substantial within-nest variation in pollen composition. This heterogeneity likely reflects flexible provisioning behavior rather than methodological noise, as maternal bees may provision different brood cells with different pollen mixtures to reduce the risk of total brood failure if a specific resource proves nutritionally deficient (Eckhardt et al., 2014). These two sampling strategies thus capture complementary aspects of foraging behavior, and future work linking within-nest variation to offspring sex allocation or survival would help clarify its adaptive significance (Wittmann et al., 2025).

The *rbcL* marker provides limited taxonomic resolution, often restricting plant identification to the genus or family level (Bell et al., 2017; Richardson et al., 2015). Although the limited geographic scale reduces the number of potential congeners, this limitation likely makes our tests for dietary partitioning conservative, because if species show distinct preferences at the family level, actual niche differentiation at the species level is likely even more pronounced. Future studies incorporating multi-locus metabarcoding or local reference libraries would improve resolution and enable finer-scale characterization of plant-bee interactions (Encinas-Viso et al., 2023; Weber et al., 2023).

### Implications for pollinator conservation in forests landscapes

Given that cavity-nesting bees rely on forests for nesting while drawing most pollen from the surrounding landscape, conservation in managed forests requires attention to both dimensions. Forest stands provide stable nesting environments whose interior conditions enhance reproductive capacity, and this nesting function has conservation value independent of whether the planted trees themselves provide sufficient floral resources. However, because planted trees alone cannot meet the pollen demands of bees throughout the nesting season, bee communities depend on phenologically complementary flowering vegetation in the surrounding matrix, including forest edges, early-successional vegetation, and shrublands (Roberts et al., 2017). Effective conservation therefore depends less on maximizing within-stand tree species richness and more on retaining or restoring diverse flowering habitats within typical bee foraging ranges of 200–500 m (Bishop et al., 2024; Zurbuchen et al., 2010). For subtropical forest plantations, preserving a mosaic of early-successional margins and shrublands is likely more effective than attempting to maximize floral diversity within plantation boundaries alone (Hall et al., 2019; Senapathi et al., 2015).

## Acknowledgments

We are grateful to the BEF–China consortium for their support, and thank Professor Bo Yang, Dr. Shan Li, and Mr. Zhi-Gao Li for their invaluable assistance. We also thank Mr. Yin-Quan Qi for his help on sample collection and Mr. Yuan Feng for his expertise in specimen identification. This work was supported by the National Natural Science Foundation of China (32470473), the Key Program of the National Natural Science Foundation of China (32330013), and the Deutsche Forschungsgemeinschaft (DFG) funded research unit MultiTroph (452861007/FOR 5281). AL was funded by the National Key Research Development Program of China (2022YFF0802300). CDZ was supported by Initiative Scientific Research Program, Institute of Zoology, Chinese Academy of Sciences (2025IOZ06). MCO was supported by the Chinese Academy of Science’s President’s International Fellowship Initiative (2026PVC0122).

